# Functional Analysis of TM6 MADS-box gene in the Octoploid Strawberry by CRISPR/Cas9 directed mutagenesis

**DOI:** 10.1101/351296

**Authors:** Carmen Martín-Pizarro, Juan Carlos Triviño, David Posé

**Affiliations:** Laboratorio de Bioquímica y Biotecnología Vegetal, Instituto de Hortofruticultura Subtropical y Mediterránea (IHSM), Universidad de Málaga-Consejo Superior de Investigaciones Científicas, Departamento de Biología Molecular y Bioquímica, Facultad de Ciencias, UMA, Málaga, Spain.; Sistemas Genómicos, Valencia, Spain.

## Abstract

The B-class of MADS-box transcription factors has been studied in many plant species, but remain functionally uncharacterized in the *Rosaceae* family. APETALA3 (AP3), a member of this class, controls the identity of petals and stamens in *Arabidopsis thaliana*. In this work, we identified two members of the AP3 lineage in the cultivated strawberry (*Fragaria* × *ananassa*): *FaAP3* and *FaTM6*. Interestingly, *FaTM6*, and not *FaAP3*, shows an expression pattern equivalent to that of *AP3* in *Arabidopsis*. Genome editing using Cluster Regularly Interspaced Short Palindromic Repeats (CRISPR)/Cas9 system is becoming a robust tool for targeted and stable mutagenesis of DNA. However, whether it can be efficiently used in an octoploid species such as *F*. × *ananassa* is not known. Here we report for the first time the application of CRISPR/Cas9 in *F*. × *ananassa* to characterize the function of FaTM6 in flower development. An exhaustive analysis by high-throughput sequencing of the *FaTM6* locus spanning the target sites showed a high efficiency genome editing already in the T0 generation. The phenotypic characterization of the mutant lines indicates that FaTM6 plays a key role in petal and especially in anther development in strawberry. in an octoploid species such as *F*. × *ananassa*, and offer new opportunities for engineering strawberry to improve traits of interest in breeding programs.

## Introduction

In the 26 years that have passed since the formulation of the classic ABC model for floral organ identity (Coen and Meyerowitz, 1991), our understanding of the molecular mechanisms controlling floral organ development has progressed significantly. The activity of the B-class proteins, APETALA3 (AP3) and PISTILLATA (PI), specifies petal and stamen identity when their activity overlaps with A-class and C-class proteins, respectively (Krizek and Meyerowitz, 1996). *AP3* and *PI* arose from an ancestral duplication event, which it is suggested that occurred before the diversification of the angiosperms (Kramer et al., 1998). Later, *AP3* experienced a second duplication before the diversification of the higher eudicots in the *AP3* lineage, resulting in two paralogous lineages, *euAP3* and *Tomato MADS box gene6 (TM6),* which differ in their C-terminal sequence motifs (Pnueli et al., 1991; Kramer et al., 1998). Although there are species that has lost one of the lineages, such as *Arabidopsis* and *Antirrhinum*, which lack *TM6*, and papaya, which lost *euAP3* instead (Causier et al., 2010), most species posses both *euAP3* and *TM6* genes, which have functionally diversified. *euAP3* genes, such as the *Arabidopsis AP3,* are mainly involved in both petal and stamen development (Jack et al., 1992). By contrast, *TM6-like* genes have a predominant role in stamens (de Martino et al., 2006; Rijpkema et al., 2006; Roque et al., 2012).

In strawberries (genus *Fragaria*), flowers differ from those of *Arabidopsis* in several aspects. The most striking difference is the presence of hundreds of independent carpels located on an enlarged stem tip, the receptacle, which expands upon carpel fertilization to generate the fleshy part of the berry, surrounded by the true fruits, the achenes (Nitsch, 1950; Hollender et al., 2011). The role of any of the homeotic genes in the ABC model of flower development has not yet been functionally determined in strawberries, so it is not yet known whether or how these genes contribute to the development of this particular type of flower.

Strawberries possess a wide range of ploidy levels, varying from diploid, such as the ancestral species, the woodland strawberry *F. vesca* (2*n* = 2*x* = 14 chromosomes), to decaploid, such as *F. iturupensis* (2*n* = 10*x* = 70); the cultivated strawberry (*F. × ananassa*) is an octoploid species (2*n* = 8*x* = 56). Recent studies have proposed that the complex origin of the *F. × ananassa* genome is the result of the hybridization of three or four different species with different levels of ploidy (Tennessen et al., 2014; Sargent et al., 2016). Recently, a virtual reference genome of *F. × ananassa* has been established after sequencing some wild relatives (Hirakawa et al., 2013), however, a whole-genome sequence for this species has not been published yet. As an alternative, the genome of the diploid *F. vesca* is commonly used as the reference genome (Shulaev et al., 2011; Tennessen et al., 2013; Tennessen et al., 2014; Li et al., 2017; Edger et al., 2018). Reverse genetics strategies employed to characterize gene function in strawberry are based on gene down-regulation via post-transcriptional gene silencing by RNA interference (RNAi) (Guidarelli and Baraldi, 2015). However, the RNAi approach has some drawbacks, such as temporary knockdown effects, unpredictable off-target influence and too much background noise (Martin and Caplen, 2007). Recent progress in genome editing methods has opened new possibilities for reverse genetics studies. In particular, the Clustered Regularly Interspaced Short Palindromic Repeat (CRISPR)/CRISPR-associated 9 endonuclease (Cas9) technology, hereafter CRISPR/Cas9, has become a very powerful tool for the acquisition of desired mutations due to its simplicity, efficiency, and stability. CRISPR/Cas9-mediated mutagenesis has been widely applied to plant research in the last few years, not only in *Arabidopsis*, (Jiang et al., 2013; Li et al., 2013; Feng et al., 2014; Gao and Zhao, 2014; Jiang et al., 2014) but also in *Rosaceae* species, such as apple (Malnoy et al., 2016; Nishitani et al., 2016), and recently, the diploid wild strawberry *F. vesca* (Zhou et al., 2018). CRISPR/Cas9 has also been used in crops with high ploidy levels such as citrus (triploid), potato, oilseed rape, cotton (tetraploids), and bread wheat (hexaploid) (Weeks, 2017). However, the functionality of this genome editing system has yet to be tested in an octoploid such as *F. × ananassa.*

In this study, we use CRISPR/Cas9 to functionally characterize the role of a homeotic gene in *F. × ananassa,* in particular, *FaTM6*, which mutation affects the development of petals, anthers, pollen grains, and, subsequently, of berries. This work demonstrates that FaTM6 plays a role equivalent to AP3 in *Arabidopsis*, and that the CRISPR/Cas9 system can be a suitable tool for functional analyses and molecular breeding in the cultivated strawberry species, which may have important economic implications.

## Results

### Identification and phylogenetic analysis of *AP3* lineage genes in *F. vesca*

To identify genes belonging to the *AP3* lineage in strawberry, we BLASTed the AP3 protein sequence from *Arabidopsis* (AtAP3) using the reference genome of the diploid *F. vesca* (cv. Hawaii 4), obtaining two genes with high homology: FvH4_2g38970 and FvH4_1g12260, sharing 55.45% and 50.43% of amino acid identity with AtAP3 respectively. To place these two genes in a phylogenetic context, we performed a phylogenetic analysis using Neighbor-Joining of AP3- and TM6-like proteins from gymnosperms to core eudicots (Fig. S1). The phylogenetic analysis shows that FvH4_2g38970 (hereafter named FveAP3) and FvH4_1g12260 (hereafter named FveTM6) belong to the *euAP3* and *TM6* lineages respectively, indicating that *F. vesca*, unlike *Arabidopsis*, contains both AP3 lineages.

### Expression analysis of *AP3* lineage genes in *F.* × *ananassa*

To further investigate the role of the *AP3* lineage genes in the cultivated strawberry, we first analyzed their expression using quantitative real-time PCR (qRT-PCR) in sepals, petals, stamens, receptacles and carpels of *F. × ananassa* flowers at stage 12 (Hollender et al., 2011). As shown in Fig. 1A, *FaTM6* is expressed in both petals and stamens, being the latest the tissue with the highest expression level. Differently, *FaAP3* is expressed mainly in receptacles, followed by carpels and petals, with very little expression in stamens and sepals (Fig. 1B). This suggests that *FaTM6*, and not *FaAP3* is the gene with the homologous function to *AP3* in *Arabidopsis*. Hence, we selected *FaTM6* to study its role in flower development by CRISPR/Cas9-mediated mutagenesis.

**Figure 1.**
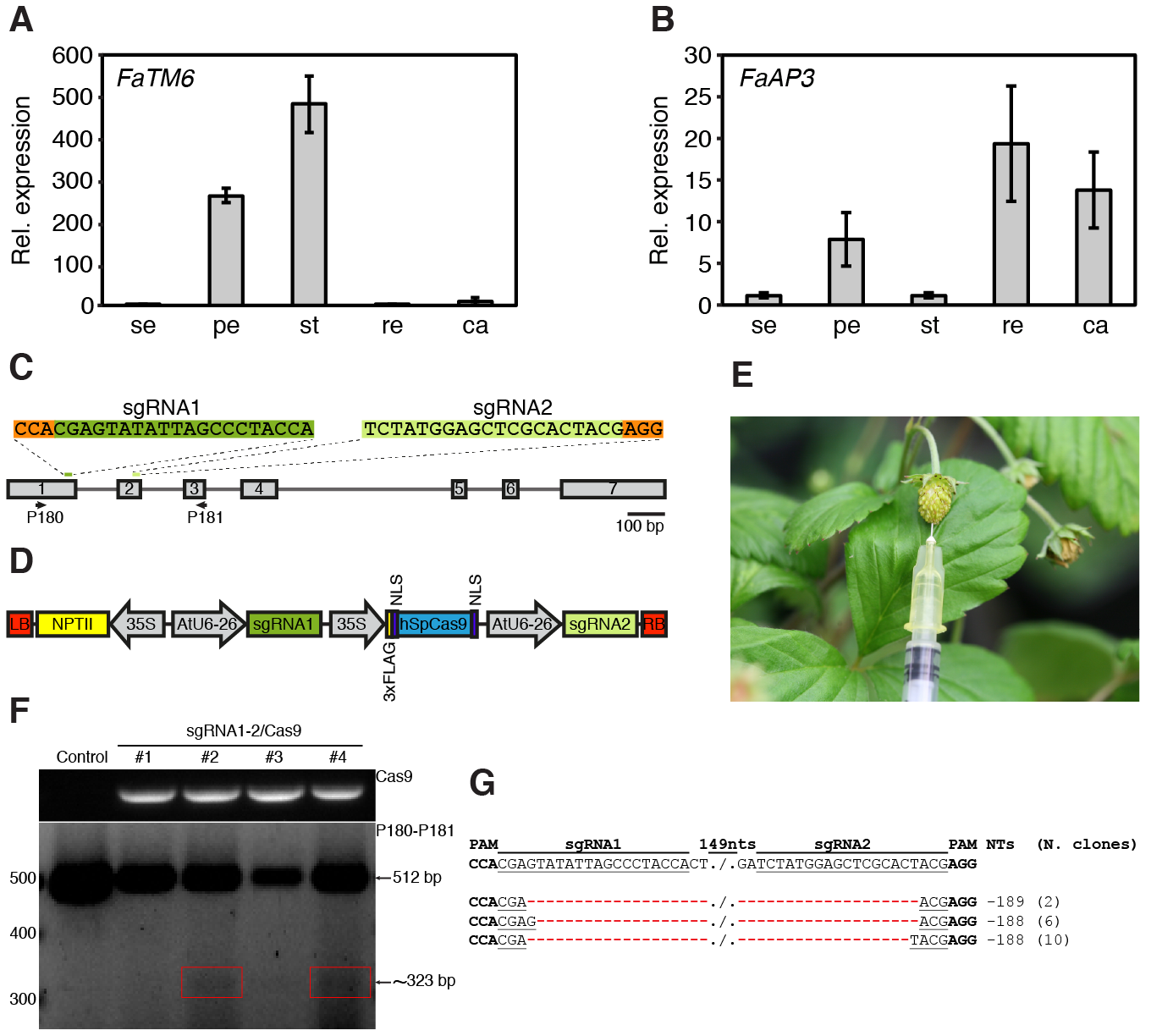
Expression of *AP3* and *TM6* genes in the cultivated strawberry (*Fragaria × ananassa)*, construct design and evaluation of CRISPR-Cas9-based editing of the wild strawberry, *Fragaria vesca*. Relative (Rel.) expression by qRT-PCR of the *F. × ananassa* (**A**) *TM6* gene (*FaTM6*) and (**B**) *AP3* gene (*FaAP3)* in sepals (se), petals (pe), stamens (st), receptacles (re) and carpels (ca) of *F. × ananassa* flowers at stage 12. Error bars denote the standard deviation (s.d.) of three biological replicates with three technical replicates each. (**C**) The *F. vesca TM6* (*FveTM6*) locus, including exons (boxes) and introns (lines). sgRNAs 1 and 2 are represented in green and PAM sequences in orange. Primers used for CRISPR/Cas9 editing characterization are indicated. (**D**) Schematic representation of the two sgRNAs and Cas9 expression cassettes in a single binary vector pCAMBIA2300 (sgRNA1-2/Cas9 vector). (**E**) Green fruit agroinfiltrated with the sgRNA1-2/Cas9 vector. (**F**) PCR to detect indels in *FveTM6*. Top panel: PCR of Cas9 in four fruits that transiently expressed the vector. Bottom panel: PCR with P180 and P181 (Table S2) showing a ~323-bp band (red squares) in fruits #2 and #4 in addition to the *wild-type* band (512 bp). (G) Alignment of sequences obtained after the purification, cloning and Sanger sequencing of the ~323 bp band from fruits #2 and #4. Fragments with 189- and 188-bp deletions resulting from simultaneous DSBs in both target sites were detected in 2 and 16 clones, respectively.

### *FaTM6* targets and off-target identification

We first searched for candidate sgRNAs to edit *FaTM6* using the available sequence of *FveTM6* from the *F. vesca* reference genome (cv. Hawaii 4). Two target sites for *FveTM6* were selected in order to generate a dual sgRNA construct that may create a large deletion and/or increase the efficiency of the mutagenesis (Belhaj et al., 2013). The sgRNAs, located within exon 1 (sgRNA1) and exon 2 (sgRNA2), were selected based on their high specificity score and minimum possible off-target activities (Fig. 1C, Table S1). The sgRNA1 spans the carboxyl-end of the M-domain and the amino-terminal of the “intervening” (I) region of the FveTM6 protein, while the sgRNA2 is located within the I region (Fig. S2). Out of the seven putative off-targets predicted, only two were located at coding sequences (CDS), containing four and five mismatches within the sgRNA1-PAM sequence respectively (Table S1). In addition, these two genes are very lowly expressed in flower organs such as petals and anthers, where *TM6* shows the highest expression in both *F. × ananassa* (Fig. S3) and *F. vesca* (Hawkins et al., 2017). Hence, we did not expect any phenotypic effect on those tissues due to a possible off-target activity.

### Identification of *FaTM6* alleles and construct design

We evaluated the suitability of the two guides designed using the reference genome for editing ability in the two genotypes used in this study (*F. vesca* cv. Reine des Vallées (RV) and *F. × ananassa* cv. Camarosa). In order to detect any possible polymorphisms to the reference genome that might affect the CRISPR/Cas9-mediated editing, we amplified and cloned the genomic regions of *TM6* spanning the two target sites (Fig. 1C, Table S2) and performed Sanger sequencing. While *F. vesca* cv. RV did not show any variation in the *TM6* sequence compared to the reference genome, five different alleles were identified in *F. × ananassa* cv. Camarosa (Fig. S4). These alleles contained indels in the first and second intron, and 4 synonymous and 8 non-synonymous SNPs within the coding region (Fig. S4). None of these alleles had polymorphisms in the region targeted by sgRNA2, but allele #5 contained a G168T substitution within the PAM-proximal region of sgRNA1 (Fig. S4), which might decrease the cleavage efficiency (Xu et al., 2017). To determine which alleles are expressed in petals and stamens, we generated cDNA from these tissues and performed high-throughput amplicon sequencing for the region spanning sgRNA1 and sgRNA2. Our data indicate that at least four of the five *FaTM6* alleles identified are expressed in both petals and stamens (Table S3). In detail, we detected alleles #3, #4, #5, and a sequence that might correspond to either allele #1 or #2, which are indistinguishable within the CDS region sequenced. Given this information and with our dual sgRNA strategy, sgRNA2 would likely ensure editing events due to the lack of polymorphisms, while sgRNA1 would allow us to assess the effect of the mismatch in allele #5 on editing efficiency. This design would also result in large deletions if both sites are cleaved by Cas9, likely producing non-functional alleles. We designed a single binary vector harboring the two sgRNAs under AtU6-26 promoters, and the Cas9 nuclease under the 35SCaMV promoter: sgRNA1-2/Cas9 (Fig. 1D).

### Functionality test of the sgRNA1-2/Cas9 vector by transient transformation of *F. vesca* fruits

Since the transformation and establishment of stable transgenic plants requires 6-9 months, we first tested the functionality of our dual sgRNA/Cas9 editing construct by transient transformation of diploid *F. vesca* cv. RV fruits. The sgRNA1-2/Cas9 vector was agroinfiltrated in the receptacle of fruits at the green stage (Fig. 1E), and genomic DNA was extracted after ten days post-infiltration. A PCR-amplification with primers spanning the target sites (Table S2) was performed in order to detect CRISPR/Cas9-mediated editing events evidenced by amplicon sizes that are different from the *wild-type* allele (Fig. 1C). The PCR results confirmed that two of the four fruits infiltrated with the sgRNA1-2/Cas9 vector showed a smaller amplicon in addition to the *wild-type* amplicon (Fig. 1F). Cloning and Sanger sequencing of the smaller amplicon from these two plants confirmed the presence of a deletion of ~190 bp between the two target sites in 18 clones, validating the functionality of the sgRNA1-2/Cas9 vector in the diploid strawberry species (Fig. 1G).

### Targeted mutagenesis of *FaTM6* in stable transgenic *F. × ananassa* plants

Next, we generated stable transgenic plants of *F. × ananassa* cv. Camarosa with the same sgRNA1-2/Cas9 vector. We obtained and micropropagated five independent lines, termed *tm6* lines, and used PCR to examine the presence of editing events as described for the transient assays (Fig. 2A; Table S2). Different amplicon patterns were obtained between the *tm6* lines and the untransformed control line, indicating the generation of various indels by the CRISPR/Cas9 complex (Fig. 2B). These results confirm the CRISPR/Cas9-mediated mutagenesis of *FaTM6* in *F. × ananassa*.

**Figure 2.**
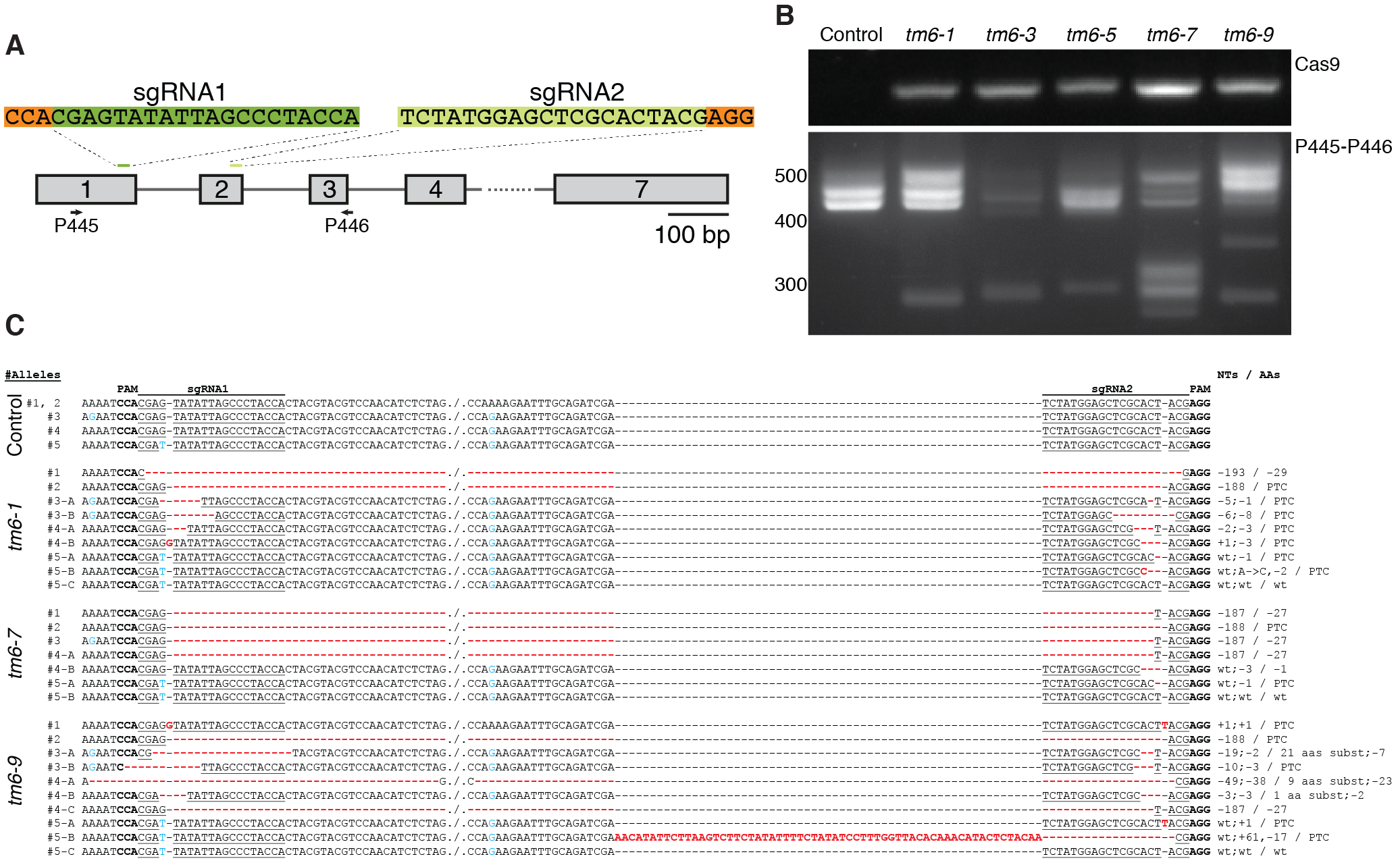
Identification of CRISPR/Cas9-induced mutations in the *F*. × *ananassa TM6* allele (*FaTM6*). (**A**) Schematic representation of the positions of the sgRNAs in the *F. vesca TM6 (FveTM6)* locus. Primers used for the analysis in agarose gel and for deep sequencing are represented (P445 and P446; Table S2). (**B**) Top panel: Identification of *Cas9* gene in transgenic *tm6* lines. Bottom panel: Detection of mutations at the *FaTM6* locus using P445 and P446 primers. (**C**) Sequence alignment obtained by high-throughput amplicon sequencing in control and *tm6* mutant lines. PAM sequences are marked in bold; sgRNAs are underlined; blue font indicates distinctive SNPs among *FaTM6* alleles; bold red font indicates mutations induced by CRISPR/Cas9-mediated editing. PTC: Premature termination codon.

In order to analyze the CRISPR/Cas9-induced mutations at the *FaTM6* locus at the molecular level, we selected three transgenic lines *tm6-1, tm6-7* and *tm6-9*, based on their amplicon patterns. Using high throughput amplicon sequencing and *de novo* assembly, nine, seven, and ten different alignment groups were obtained for *tm6-1, tm6-* 7, and *tm6-9* respectively (Fig. 2C; Table S3). Alleles #1 showed a single editing event in all three lines, while allele #2 had the same deletion in all of them (Fig. 2C). However, different editing events were obtained for alleles #3, #4 and #5 within each transgenic line, except for *tm6-7*, which had only one modification in allele #3 (Fig. 2C). As expected, most of the CRISPR/Cas9-induced mutations occurred downstream of the PAM sequences, although a deletion including the whole PAM-sgRNA1 region was also observed in *tm6-9* (Fig. 2C).

All sequences obtained for alleles #1 through #4 showed mutations within the region targeted by sgRNA1 (Fig. 2C). However, no editing was observed in this target site in allele #5, most likely due to the mismatch present in the sgRNA1 seed sequence (Fig. 2C). For the region targeted by sgRNA2, the sequencing analysis showed that all five alleles were edited, although *wild-type* sequences were also detected for this target only in allele #5 (Fig. 2C).

As expected from the amplicon pattern obtained for these three lines, all of them contained large deletions generated by the simultaneous double-strand breaks (DSBs) in both target sites. These deletions of 187 and 193 nts resulted in a deletion of 27 or 29 amino acids (aas), respectively, or in the generation of a premature stop codon (™188 nts) (Fig. 2C and Fig. S5). Most of the editing events generated frameshift mutations, especially in the *tm6-1* line, which resulted in the production of seven truncated proteins (#2, #3A, #3B, #4A, #4B, #5A and #5B) out of the nine allelic variants detected (Fig. S5). In addition to the generation of these truncated proteins, shorter amino acid deletions and substitutions were also obtained in *tm6-7* and *tm6-9* (Fig. S5).

### FaTM6 plays a key role in anther development

To determine the function of *FaTM6*, we analyzed the flower phenotype of the *tm6-1, tm6-7* and *tm6-9* lines. At pre-anthesis stage, petals in the mutant lines were shorter and greenish compared to that of the control flowers (Fig. 3A-B). More severe defects were observed in the anthers, which were smaller and darker than that of the wild-type (Fig. 3C). A more detailed anatomical analysis of the anthers at the dehiscence stage and that of the pollen grains were performed by Scanning Electron Microscopy (SEM). *Wild-type* anthers displayed the typical four-lobed structure, with a very well defined epidermal layer, and with pollen grains visible at the stomium rupture site (Fig. 3D). However, the anthers of *tm6* mutant lines displayed morphological differences, showing clear defects in the epidermal cell layer and a reduced number of pollen grains at the stomium (Fig.3D). This apparent difference in pollen content was quantified, resulting in a 10-fold reduction in *tm6-1* and *tm6-7*, and a 50-fold reduction in *tm6-9* compared to the control (Fig. S6). Moreover, most of the pollen grains from the mutant lines showed aberrant and collapsed structures (Fig. 3E).

**Figure 3.**
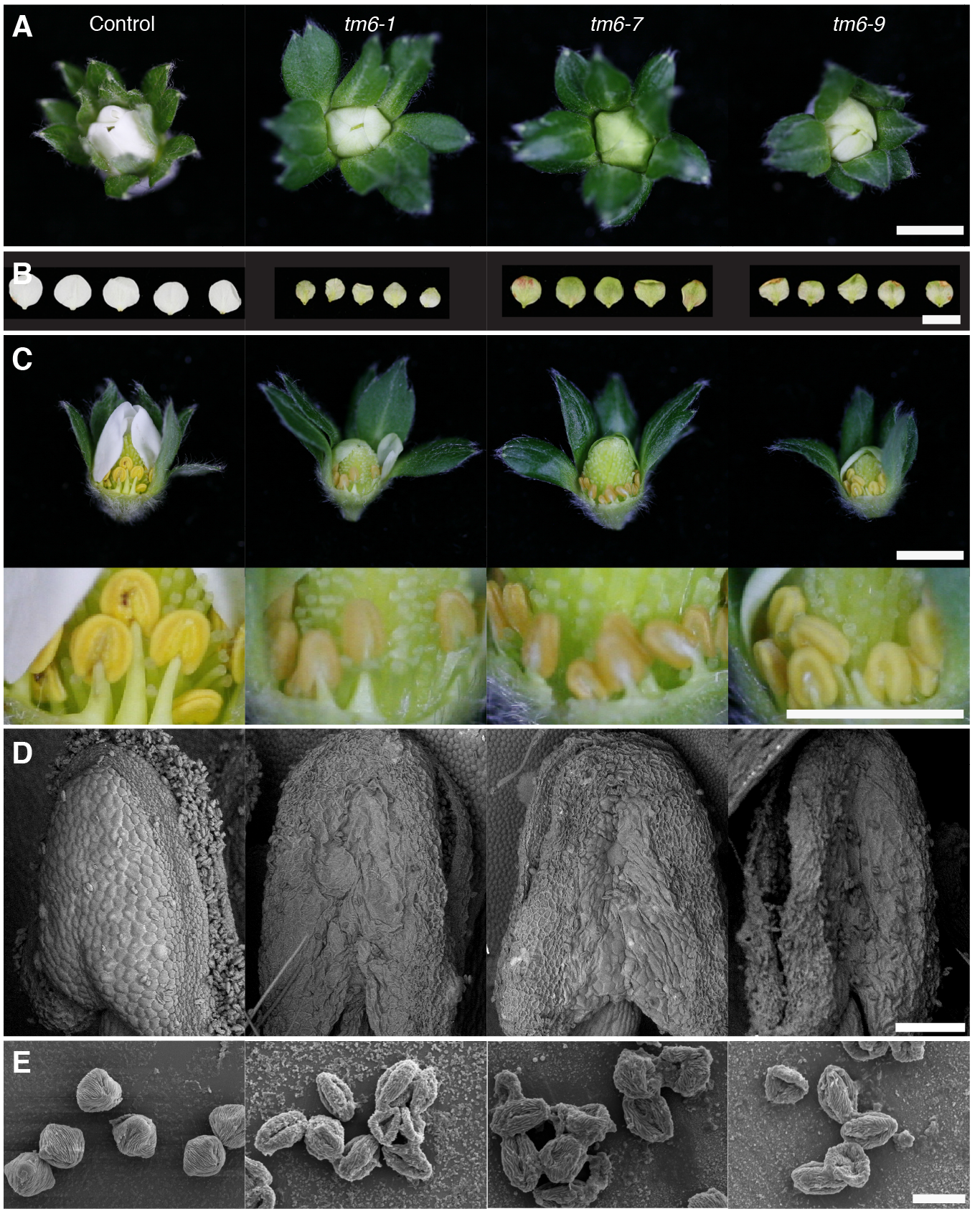
Phenotypic effects of mutations in *F. × ananassa TM6 (FaTM6)* in flowers. (**A**) Flowers of control and three independent *tm6* lines at the pre-anthesis stage. (**B**) Petals of *tm6* lines appear smaller and greenish. (**C**) Top panel: flowers at pre-anthesis with some petals removed. Bottom panel: higher magnification to show details of the morphology of the stamens. (**D-E**), Scanning electron microscopy (SEM) of the structure of the anthers at the dehiscence stage (**D**) and pollen grains (**E**). Scale bars: (**A-C**): 1 cm; (**D**): 200 μm; (**E**): 20 μm.

Since ovule fertilization and proper development of embryos and achenes are necessary for normal receptacle development (Nitsch, 1950), we assessed the fruit formation in *tm6* lines compared with emasculated controls. The emasculation of WT flowers caused a complete abortion in the receptacle development (Fig. 4A). Consistent with the impaired pollen grain formation, the *tm6* mutant lines also showed arrested development of the receptacles (Fig. 4A, B). Nevertheless, a few fruits (5.7 and 2.2% in *tm6-7* and *tm6-9* respectively) showed a local enlargement of the receptacle around some achenes, indicating that some residual pollination took place (Fig. S7).

**Figure 4.**
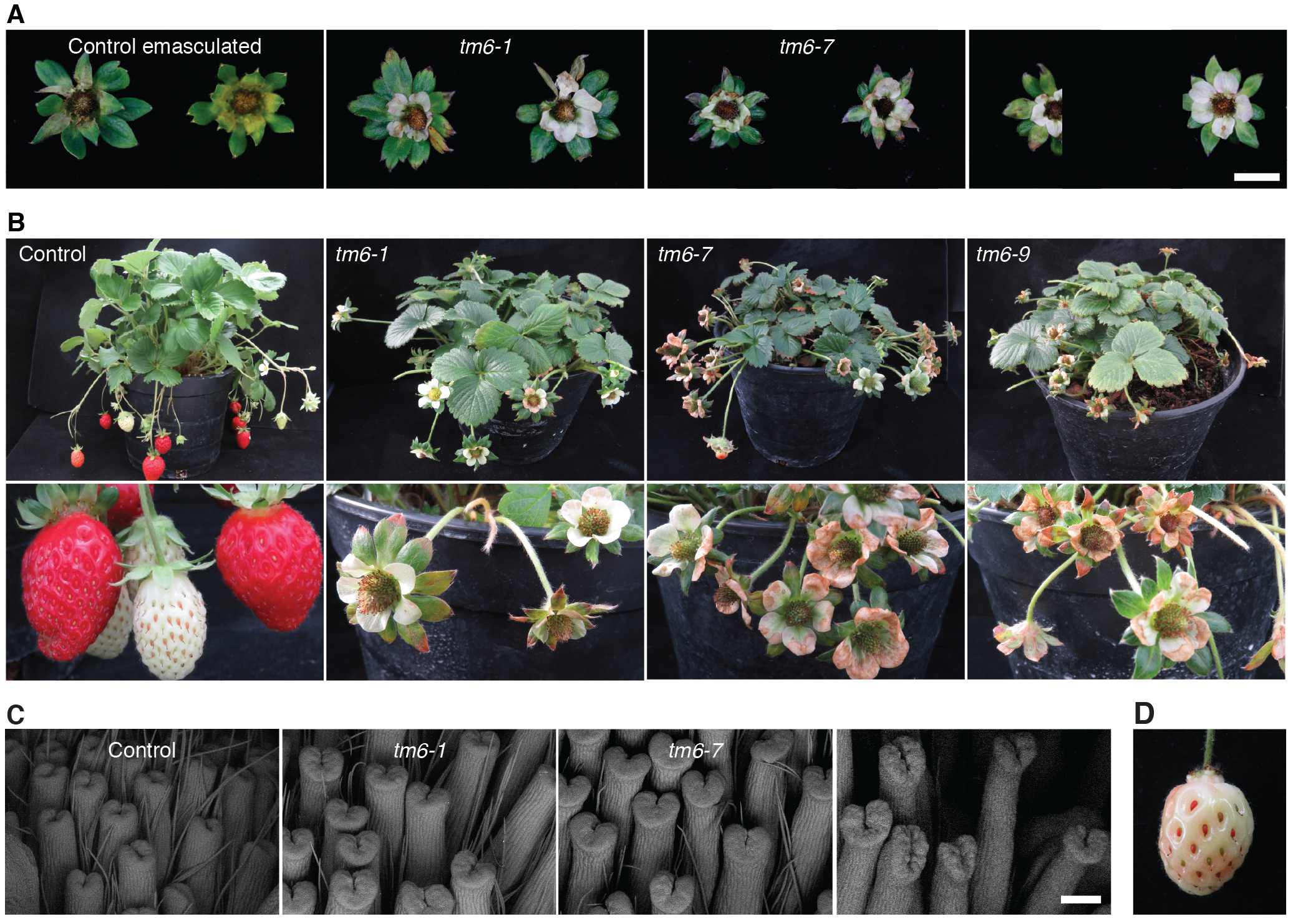
Phenotypic effect of mutations in *F. × ananassa TM6 (FaTM6)* in fruits and complementation experiment. (**A**) *Wild-type* flowers emasculated at the pre-anthesis stage phenocopy aborted flowers in *tm6* mutant lines. (**B**) Top panels: adult plants of control and *tm6* mutant lines. Bottom panels: control plant develops *wild-type* berries, but *tm6* flowers abort. (**C**) Scanning electron microscopy (SEM) of the structure of carpels at pre-anthesis stage. (**D**) Fruit developed from a *tm6-7* flower emasculated and pollinated with *wild-type* pollen. Scale bar: (A): 1 cm; (C): 200 μm.

Consistent with the lack of *FaTM6* expression in carpels, these organs showed normal development (Fig. 4C). In order to confirm the carpel viability in the mutant lines, we pollinated carpels of the *tm6-7* line using WT pollen. As shown in Fig. 4D a WT receptacle was fully developed, indicating that the lack of pollination is the responsible for the aborted fruit phenotype in the *tm6* lines. These results taken together indicate an essential role of *FaTM6* in anther and pollen formation during flower development of *F. × ananassa*.

## Discussion

### FaTM6 is involved in petal and stamen development

Here, we report that strawberry, unlike *Arabidopsis thaliana*, has maintained both a *euAP3* and a *TM6* gene. While FveAP3 contains the euAP3 motif in the C-terminal domain of the protein, FveTM6 posses a motif more similar to those of the paleoAP3 genes (Fig. S2) (Kramer et al., 1998). A transcriptome study during floral development of *F. vesca* reported that *FveTM6* (gene14896-v1.0-hybrid) is strongly expressed in anthers (Hollender et al., 2014), consistent with our result in the octoploid strawberry. However, in that work, *FveTM6* was mis-annotated as *FveAP3*, since our phylogenetic analysis shows that this gene belongs to the *TM6* lineage instead (Fig. S1). Moreover, expression analysis of *FaAP3* and *FaTM6* in flowers of *F. × ananassa* have shown that *FaTM6* and not *FaAP3* is the gene with the typical B-class type expression pattern (Fig. 1A-B), which is consistent with previous studies in *Rosaceae* (Hibino et al., 2006). Sequencing analyses of the region spanning the CRISPR target sites for *TM6* indicate that the diploid *F. vesca* cultivar used in this study is homozygous at this locus. However, the octoploid strawberry *F. × ananassa* cv. Camarosa showed high heterozygosity at the *FaTM6* locus, consistent with the genetically complex genome of this species. At least five alleles of *FaTM6* among the four pairs of homoeologous chromosomes were detected. However, we cannot discard an even more complex scenario with more *FaTM6* alleles due to the possible presence of additional polymorphisms outside the region covered in this study. Furthermore, deep sequencing of cDNA from petals and sepals showed that at least four of the five *FaTM6* are expressed in both organs, being alleles #1 and #2 indistinguishable within the CDS region sequenced (Table S3).

The role of *euAP3* genes in petal and stamen specification has been well established (Jack et al., 1992; Schwarz-Sommer et al., 1992; de Martino et al., 2006; Roque et al., 2012). However, there have been fewer reports on the role of TM6-like TFs, which play a more important role in stamen identity (Rijpkema et al., 2006; Roque et al., 2012). Our results are also consistent with a predominant role of FaTM6 in stamen development since the anthers in the *tm6* mutant lines were severely affected, and they showed a drastic reduction in pollen content and viability (Fig. 3C-E, Fig. S6), while the petals of the *tm6* mutants showed more modest defects in overall size and color (Fig. 3B). This is consistent with the phenotype reported in TM6i (RNAi) lines of tomato, which developed smaller petals that were attributed to a decrease in cell proliferation (de Martino et al., 2006). Similarly, *PTD*, the *Populus trichocarpa TM6* ortholog, has been postulated to play a role in regulating cell proliferation (Sheppard et al., 2000) Auxin transport from the fertilized carpels to the floral receptacle is essential for the latter to grow into a fleshy and edible fruit (Nitsch, 1950; Kang et al., 2013). The high percentage of fruit abortions obtained in the *tm6* mutant lines, which phenocopy emasculated WT flowers, supports the tight coupling of flower and fruit development (Fig. S7, Fig. 4A-B). The presence of a small number of aberrant fruits with enlarged portions of receptacle around developed achenes (Fig. S7) indicates that some residual viable pollen is formed. *tm6* mutants developed anatomically normal pistils (Fig. 4C), consistent with the lack of *FaTM6* expression in this organ (Fig. 1A). In fact, carpels in *tm6* lines are functional since fruit development was restored using WT pollen. All of these findings indicate that the defect in pollen formation and not in gynoecium development causes the fruit abortions in the *tm6* lines.

### CRISPR/Cas9 is an efficient tool for gene functional analysis in the octoploid strawberry

In this study, we have designed a dual sgRNA system that has efficiently edited the *FaTM6* gene in the octoploid strawberry. Moreover, we have developed a quick and easy validation *in vivo* of the sgRNA efficiency performing a transient assay in fruits of the diploid *F. vesca*.

A deep sequence analysis in three independent transgenic CRISPR lines in *F. × ananassa* showed a high efficiency of sgRNA1, which drove the editing of 4 of the 5 *FaTM6* alleles in all lines examined (Fig. 2C, Table S3). Only allele #5 remained totally unedited at this target site, likely because of the mismatch in the seed sequence, which has been reported to decrease the cleavage efficiency (Xu et al., 2017). Although Cas9 can still tolerate mismatches within the target site (Hsu et al., 2013; Braatz et al., 2017), our results indicate that the mismatch in the seed sequence of allele #5 totally prevented the Cas9 activity (Fig. 2C, Table S3). None of the alleles had mismatches with sgRNA2, and this led to the editing of all the five *FaTM6* alleles. Interestingly, an unedited variant was also detected with a low prevalence in each line only for allele #5 (#5-C, #5-B and #5-C in *tm6-1, tm6-7* and *tm6-9* respectively) (Table S3). It is possible that the SNP present at the sgRNA1 target site for allele #5 also affected editing at the sgRNA2 site. All of these results support that a preliminary sequence analysis is an essential step to optimize the efficiency of the CRISPR/Cas9 system in a polyploid and highly heterozygous species such as *F. × ananassa*, especially when a reference genome sequence is not available. Moreover, when designing CRISPR/Cas9 experiments to edit a polymorphic locus of *F. × ananassa*, we recommend designing constructs containing multiple sgRNAs against the different allelic variants.

Despite the octoploid nature of strawberry, our analysis indicates that *tm6-1* and *tm6-9* lines contain more than eight allelic variants, indicating that these two lines are chimeras. These genetic mosaics are common in CRISPR/Cas9 T0 generations of plants obtained from somatic tissue due to the activity of Cas9 during later stages of shoot development (Liu et al., 2017). Due to the high heterozygosity of the cultivated strawberry, breeding lines can be only maintained and propagated clonally by runners. Therefore, even though a full knock-out was not achieved in the T0 generation, continuous clonal propagation of the transgenic lines containing the CRISPR/Cas9 may enable the eventual mutation of all eight homeologs.

In summary, we have characterized the role of a homeotic gene in strawberry, *FaTM6*, for the first time. We have shown that it is primarily involved in anther development, but also has a role in petal formation. Therefore, we propose that CRISPR/Cas9 is a powerful tool for gene functional studies in the commercial strawberry that can overcome the drawbacks of the RNAi such as instability and unpredictable off-targets. Moreover, we show that genome-editing is a feasible approach that can be used in the future to generate lines with agronomic traits of interest in *F. × ananassa* despite the high ploidy of this species.

## Methods

### Alignment and Phylogenetic Tree of AP3- and TM6-like proteins

The *A. thaliana* AP3 protein sequence was BLASTed against translated protein sequences of the strawberry genome (v4.0.a1) (Edger et al., 2018) at the Genome Database for Rosaceae server (https://www.rosaceae.org/) to obtain *Fragaria vesca* TM6 (FveTM6) and AP3 (FveAP3) protein sequences. Multiple sequence alignment of AP3-and TM6-like proteins were performed using MUSCLE with the SeaView version 4 program (Gouy et al., 2010). The phylogenetic tree was inferred by the Neighbor-Joining method. A total of 1000 bootstrap pseudo-replicates were used to estimate reliability of internal nodes. Evolutionary distances were computed using the Poisson correction method. The tree was rooted using four PI-like sequences: At-PI (*A. thaliana*), TPI (*S. lycopersicum*), FvePI-1 and FvePI-2 (*F. vesca*). Tree inference was performed using MEGA version 7 (Kumar et al., 2016). The dataset comprised 68 previously reported *AP3-* and TM6-like genes from gymnosperms, monocots, basal angiosperms, basal eudicots and core eudicots obtained from GenBank. All sequences used in this analysis, with their GenBank Accession numbers and respective species, are listed in the Accession numbers section.

### Plant material, transient and stable transformation

F. *vesca* (cv. Reine des Vallées) and *F. × ananassa* Duch. (cv. Camarosa) plants were grown and maintained under green house conditions (IHSM, Málaga, Spain). Transient expression of the sgRNA1-2/Cas9 binary vector was performed by infiltration of a suspension of *Agrobacterium tumefaciens* (strain AGL-0) into fruits at the green stage of development of *F. vesca* as previously described (Hoffmann et al., 2006). Fruits were collected ten days after the infiltration when they reached the red stage. For stable transformation of *F. × ananassa* cv. Camarosa, plants were micropropagated in N30K medium supplemented with 2.20 μM Kinetin. Transformation was performed according to the protocol described by Barceló and colleagues (Barceló et al., 1998). Leaf discs were transformed with *Agrobacterium* (strain LBA4404) carrying a pCAMBIA2300 plasmid that contained the kanamycin resistance gene *nptII* and the Cas9-sgRNA cassette. Regenerated shoots were selected in the same medium supplemented with 50 mg.1^−1^ kanamycin and 500 mg.l^−1^ carbenicillin. Resistant plants were transferred to the green house after 20-30 weeks post-transformation.

### Design of sgRNAs and vector construct

Genomic sequence of *FveTM6* (FvH4_1g12260), previously annotated as gene14896-v1.0-hybrid, was obtained from the reference genome of Shulaev et al., 2011. Single-guide RNAs were designed using the ATUM CRISPR/gRNA tool(https://www.atum.bio/eCommerce/cas9/input) with *FveTM6* CDS as the input sequence. We performed a BLAST search of *FveTM6* CDS against *F. vesca* reference genome to select those candidate sgRNAs that were specific to the target gene. Two sgRNAs located in exon 1 and exon 2, and separated by 198 bp (from PAM to PAM) were selected. CRISPOR web-tool (http://crispor.org) (Haeussler et al., 2016) was used to validate the quality of the selected sgRNAs and to identify putative off-targets.

Two different vectors were used to generate the final sgRNA1-2/Cas9 construct: pAtU6:sgRNA and 35S:hSpCas9 (Mao et al., 2013). We cloned both sgRNA1 and sgRNA2 into pAtU6:sgRNA vectors using BbsI. pAtU6:sgRNA1 was cloned into 35S:hSpCas9 vector using Acc65I and SalI. pAtU6:sgRNA2 was amplified to include Cfr9 and XbaI restriction sites and cloned it into the pAtU6:sgRNA1/35S:hSpCas9 vector, generating a construct with the Cas9 and the two sgRNAs cassettes. The final binary vector was obtained cloning the sgRNA1-2/Cas9 cassettes into pCAMBIA2300 using KpnI and XbaI sites. The binary vector was introduced into *Agrobacterium tumefaciens* strain AGL-0 for transient expression and into LBA4404 for stable transformation.

### Mutation identification

Genomic DNA was isolated using the CTAB method from fruits of the transiently transformed plants, and leaves from stably transformed plants. The presence of the transformation cassette was tested by PCR using primers P248 and P249 to amplify Cas9 (Table S2). CRISPR/Cas9-mediated indels were detected by agarose gel electrophoresis after PCR using primers flanking both sgRNAs (P180/P181 for transient assay, and P445/P446 for stable lines; Table S2). For transient expression experiments, the PCR amplicons were cloned into pGEMT-Easy vector system (Promega, Madison, USA) and transformed into *E. coli* DH5α. Single colonies were picked to identify mutations by Sanger sequencing.

### Amplicon sequencing and sequence analysis

cDNA from petals and stamens was amplified using P445 and P475 (Table S2), and genomic DNA from leaves was amplified using P445 and P446 (Fig. 2; Table S2) for high-throughput amplicon sequencing. Resulting amplicons were reamplified for indexing. Libraries were purified with Agencourt^®^ AMPure^®^ XP beads (Beckman Coulter) using the manufacturer’s recommendations, and their quality was verified using a TapeStation 4200 HS DNA (Agilent). Libraries were quantified using real time PCR. The libraries were pooled in equimolar ratios before paired-end sequencing (2 × 250 cycles) using a MiSeq system (Illumina).

For the sequence analysis, the paired-end reads were collapsed using the FLASH algorithm developed by (Magoč and Salzberg, 2011), with a quality filter of 33 in Phred scale. Then, the resulting clusters were transformed to FASTA format using a custom python script. 35 nucleotides in the 5′ and 3′ positions were trimmed using Trimmomatic in order to reduce noise (Bolger et al., 2014). Custom python scripts were designed for sequence identification and quantification. For the quantification process, no similarity threshold for sequence clusters was applied. Sequences with at least 1% of prevalence were selected as possible allelic variants. However, PCR bias hid some alleles. In these rare cases, a fingerprint strategy based in exclusive SNPs for the missing allele that allowed variations among the target sites was designed using custom python scripts.

### Phenotypic analyses

For control plants and *tm6* mutant lines, flowers at the pre-anthesis stage were marked so that all phenotypic analyses could be performed at the same developmental stage. SEM visualization of stamens was performed using flowers at 2 days post-anthesis, and that of carpels at the pre-anthesis stage. Stamens and carpels were visualized without processing using a JEOL JSM-6490LV electron microscope under low vacuum conditions (30 MPa). To analyze the pollen morphology, anthers at the dehiscence stage were incubated in absolute ethanol for 3 hours, air-dried, and coated with gold in a sputtering Quorum Q150R ES. Gold-coated pollen grains were examined using a MEB JEOL 840 microscope.

### Quantification of pollen grains

To quantify the pollen content in control and *tm6* mutant lines, three flowers per genotype were collected at 2 days post-anthesis. Anthers were removed and incubated with 10% sucrose and 1% acetocarmine for the staining of viable pollen. Pollen grains were quantified using a Neubauer chamber under a stereomicroscope Multizoom AZ-100 (Nikon).

### Emasculation and cross-pollination

Emasculation of flowers was performed by removing all of the stamens in control flowers at the pre-anthesis stage. In order to avoid cross-pollination, the emasculated flowers were covered with cotton. To analyze the functionality of the carpels in *tm6* lines, *tm6-7* flowers were pollinated using *wild-type* pollen. In detail, the anthers of the mutant flowers were removed manually at pre-anthesis to avoid any possible self-fertilization. Then, a small paintbrush was used to pollinate the *tm6* stigmas with *wild-type* pollen. In order to avoid cross-contamination, cross-pollinated flowers were covered with cotton.

### Accession numbers

Most of the protein sequences were obtained from GenBank: MASAKO B3 (*Rosa rugosa*; AB055966), PaTM6 (*Prunus avium*; AB763909), MdMADS13 (*Malus × domestica*; AJ251116), MdTM6 (*M. × domestica*; AB081093), HmTM6 (*Hydrangea macrophylla*; AF230703), GDEF1 (*Gerbera hybrida*; AJ009724), VvTM6 (*Vitis vinifera*; DQ979341), BalTM6 (*Balanophora fungosa*; JQ613232), PhTM6 (*Petunia × hybrida*; DQ539417), LeTM6 (*Solanum lycopersicum*; X60759), NbTM6 (*Nicotiana benthamiana*; AY577817), PtAP3-2 (*Pachysandra terminalis*; AF052871), PtAP3-1 (*P. terminalis*; AF052870), Gu.ti. AP3-5 (*Gunnera tinctoria*; AY337757), Gu.ti. AP3-4 (*G. tinctoria*; AY337756), FavAP31.1 (*Fragaria × ananassa*; AY429427), MASAKO euB3 (R. *rugosa*; AB099875), GDEF2 (*G. hybrida*; AJ009725), HpDEF2 (*Hieracium piloselloides*; AF180365), HpDEF1 (*H. piloselloides*; AF180364), AtAP3 (*Arabidopsis thaliana*; AF115814), CMB2 (*Dianthus caryophyllus*; L40405), SLM3 (*Silene latifolia*; X80490), RAD2 (*Rumex acetosa*; X89108), RAD1 (*R. acetosa*; X89113), JrAP3 (*Juglans regia*; AJ313089), RfAP3-2 (*Ranunculus ficaria*; AF130870), RfAP3-1 (*R. ficaria*; AF052854), HmAP3 (*H. macrophylla*; AF230702), NMH7 (*Medicago sativa*; L41727), VvAP3 (*V. vinifera*; EF418603), CitMADS8 (*Citrus unshui*; AB218614), DEF (*Antirrhinum majus*; X52023), NtDEF (*Nicotiana tabacum*; X96428), PhDEF (*P*. × *hybrida*; DQ539416), StDEF (*Solanum tuberosum*; X67511), TAP3 (*S. lycopersicum*; DQ674532), LeAP3 (S. *lycopersicum*; AF052868), RfAP3-1 (*R. ficaria*; AF052854), RbAP3-1 (*Ranunculus bulbosus*; AF052876), RfAP3-2 (*R. ficaria*; AF130870), RbAP3-2 (*R. bulbosus*; AF130869), PnAP3-1 (*Papaver nudiculae*; AF052873), PapsAP3-1 (*Papaver somniferum*; EF071993), PcAP3 (*Papaver californicum*; AF052872), LtAP3 (*Liriodendron tulipifera*; AF052878), MpMADS7 (*Magnolia praecocissima*; AB050649), Pe.am.AP3 (*Persea americana*; AY337748), CfAP3-1 (*Calycanthus floridus*; AF230699), CfAP3-2 (C. *floridus*; AF230700), PeMADS2 (*Phalaenopsis equestris*; AY378149), PeMADS5 (*P. equestris*; AY378148), OsMADS16 (*Oryza sativa*; AF077760), SILKY 1 (*Zea mays*; AF181479), PeMADS4 (*P. equestris*; AY378147), LMADS1 (*Lilium longiflorum*; AF503913), LRDEF (*Lilium regale*; AB071378), CryMADS1 (*Cryptomeria japonica*; AF097746), CryMADS2 (C. *japonica*; AF097747), GnegGGM2 (*Gnetum gnemon*; AJ132208), GnegGGM13 (*G. gnemon*; AJ132219), DAL13-1 (*Picea abies*; AF158543), PrDGL (*Pinus radiata*; AF120097), GnegGGM15 (G. *gnemon*; AJ251555), TPI (S. *lycopersicum*; DQ674531) FveTM6 (*Fragaria vesca*; FvH4_1g12260), FveAP3 (*F. vesca*; FvH4_2g38970). AtPI (*A. thaliana*; At5g20240), FvePI-1 (*F. vesca*; FvH4_2g27860.1), FvePI-2 (*F. vesca*; FvH4_2g278270.1).

## Author Contributions

C.M-P. and D.P. planned, performed and analyzed the experiments and wrote the manuscript. J.C.T. analyzed the high-throughput sequencing data. D.P. supervised the experiments and the high-throughput sequencing data analyses. All authors read and approved the final manuscript.

## Acknowledgments

We thank Beth Rowan and Catharina Merchante for their suggestions to improve the manuscript, and especially to Miguel Ángel Botella and Victoriano Valpuesta for helpful discussions. José Sánchez-Sevilla, Alicia Esteban and José Duarte for technical assistance, and Jian-Kang Zhu for the CRISPR/Cas9 vectors. This work was supported by the Grant ERC-2014-StG 638134 (European Research Council) and the Ramón y Cajal program RYC 2013-1269 (MINECO-Universidad de Málaga, Spain.

---

**Supplemental Figure 1.**
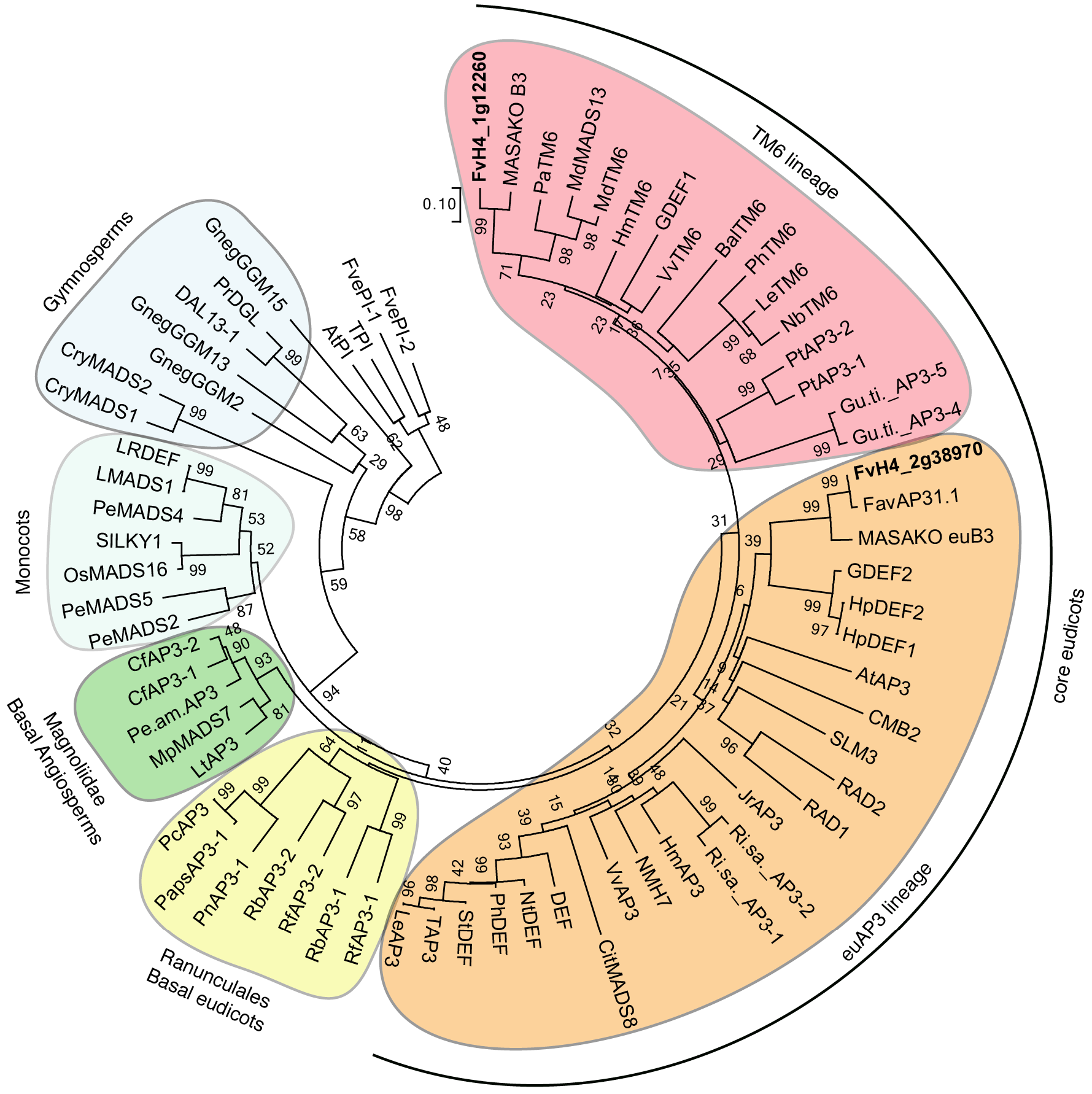
Neighbor-Joining Analysis of TM6 and euAP3 lineage proteins. Representative AP3 lineage proteins from core eudicots to gymnosperms were included in the analysis. The two AP3-like proteins from *Fragaria vesca* (FvH4_1g12260 and FvH4_2g38970) are represented in bold types. Four *PISTILLATA* genes were used as outgroup. Numbers next to the nodes are bootstrap values from 1000 pseudo-replicates. The protein sequences were obtained from GenBank (see Accesion numbers section).

**Supplemental Figure 2.**
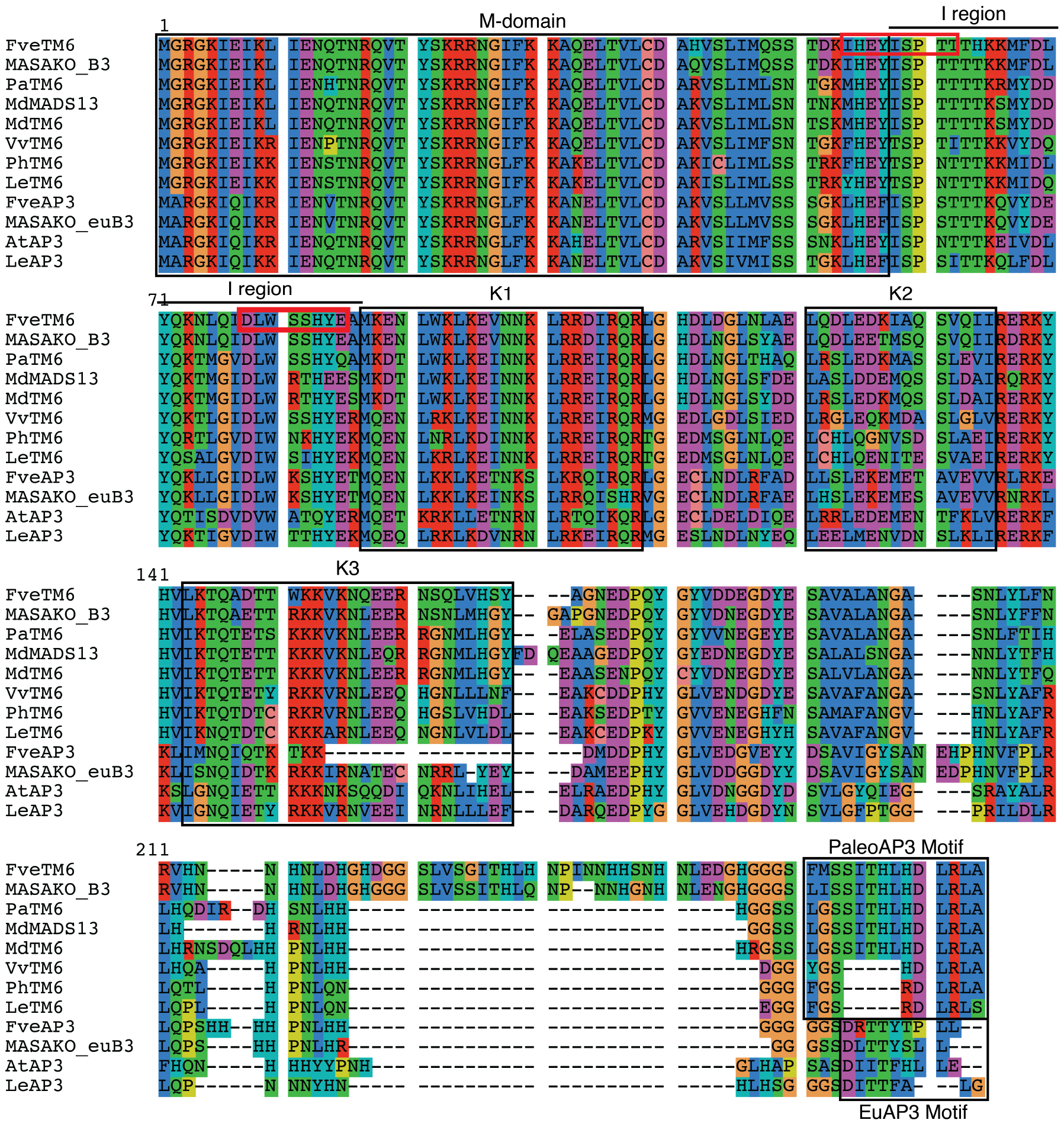
Alignment of AP3-and TM6-like proteins. Eight TM6-and four AP3-like proteins were selected for the alignment. The M-and K-domain characteristics of MIKC-type MADS transcription factors are boxed. PaleoAP3 and EuAP3 motives are located at the carboxyl end of the TM6-and AP3-like proteins. Red squares mark the region where the sgRNAs were designed for *F. vesca TM6 (FveTM6*). sgRNA1 is located spanning the M-domain and the I region. sgRNA2 is located at the I region. FveTM6 and FveAP3 (*F. vesca*), MASAKO B3 and MASAKO euB3 (*Rosa rugosa*), PaTM6 (*Prunus avium*), MdMADS13 and MdTM6 (*Malus × domestica*), VvTM6 (*Vitis vinifera*), PhTM6 (*Petunia × hybrid*), LeTM6 and LeAP3 (*Solanum lycopersicum*), AtAP3 (*Arabidopsis thaliana*). The protein sequences were obtained from GenBank (see Accession numbers section).

**Supplemental Figure 3.**
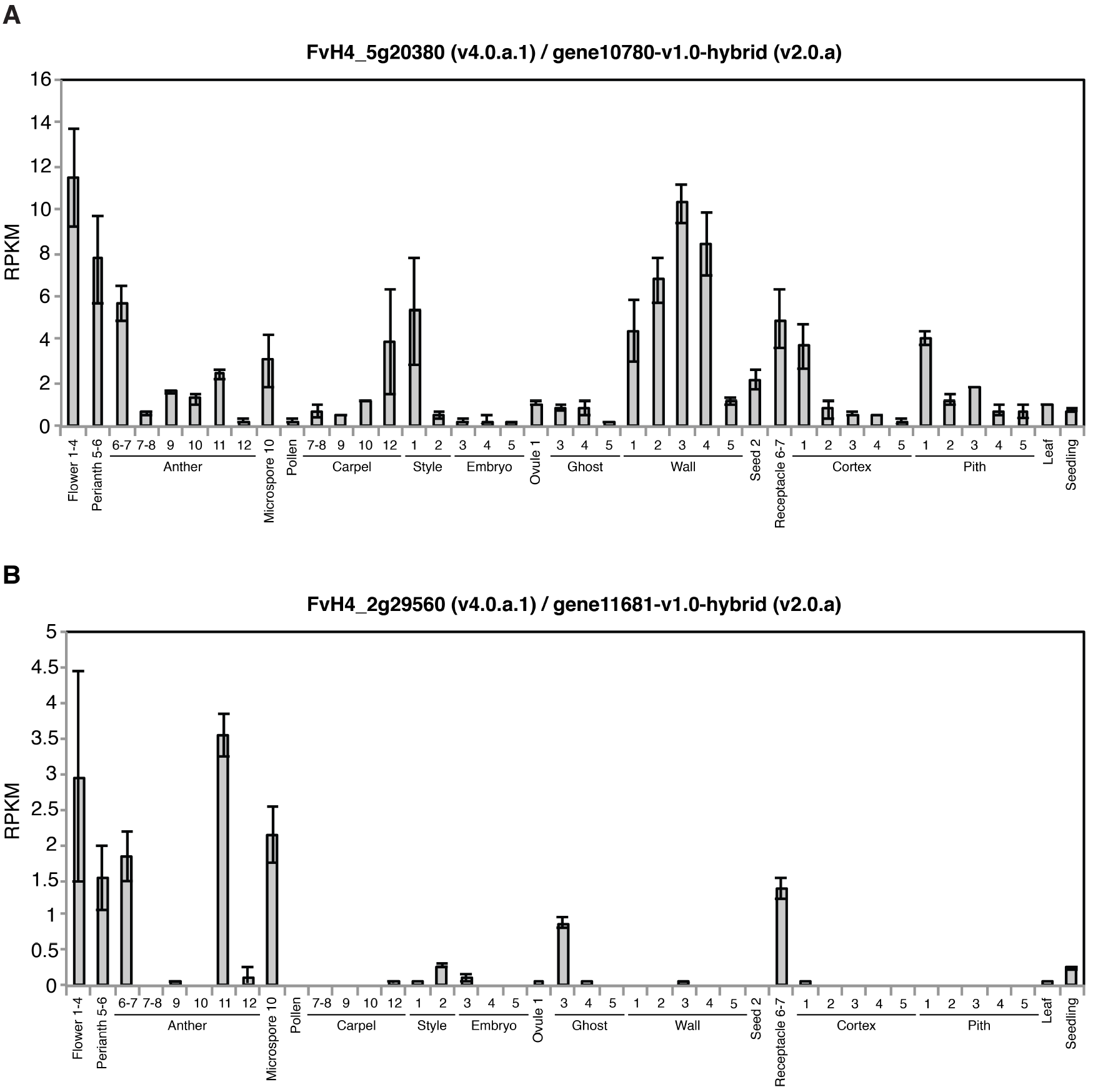
Expression analysis of two putative off-targets. Expression of FvH4_5g20380 and FvH4_2g29560 was analyzed using the eFP browser for *F. vesca* (Hawkins et al., 2017). Expression data from the flower and fruit stages were obtained from Hollender et al., 2014 and Kang et al., 2013 respectively. All stage numbering follows Hollender et al., 2011.

**Supplemental Figure 4.**
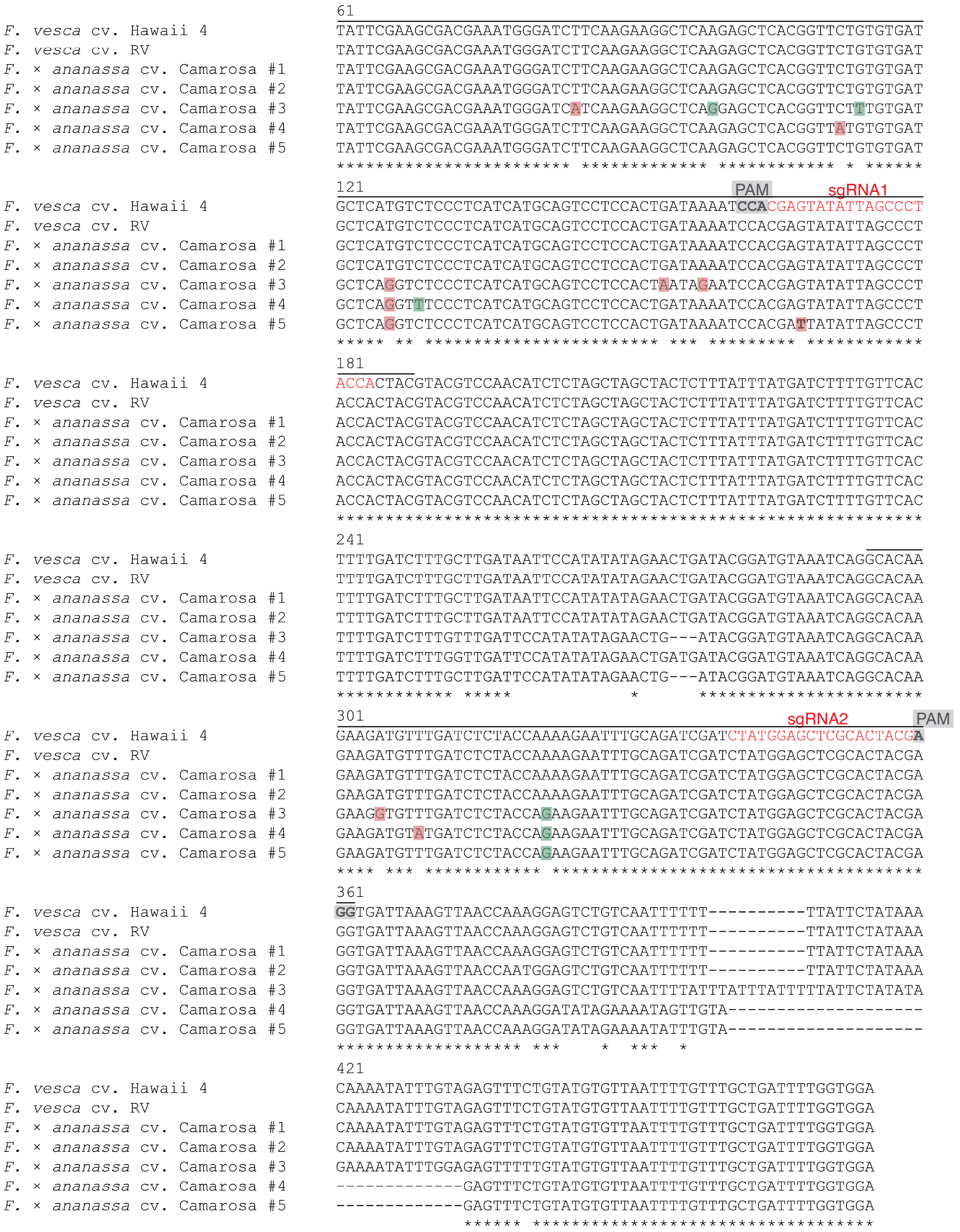
Alignment of *TM6* sequences from *F. vesca* and *F. × ananassa*. PCR flanking the two target sites (primers P180 and P181; Supplemental Table 3) for *TM6* was performed, purified, cloned and sequenced by the Sanger method for *F. vesca* cv. Hawaii 4, *F. vesca* cv. Reine des Vallées (RV), and *F. × ananassa* cv. Camarosa. The aligned region spans from the position 61 after the start codon, to the nucleotide 475, based on the *TM6* sequence in *F. vesca*. Exons are delimited with a black line; red font: sgRNAs; grey background: PAM; green background: synonymous polymorphisms; red background: non-synonymous polymorphisms; asterisks: conserved nucleotides.

**Supplemental Figure 5.**
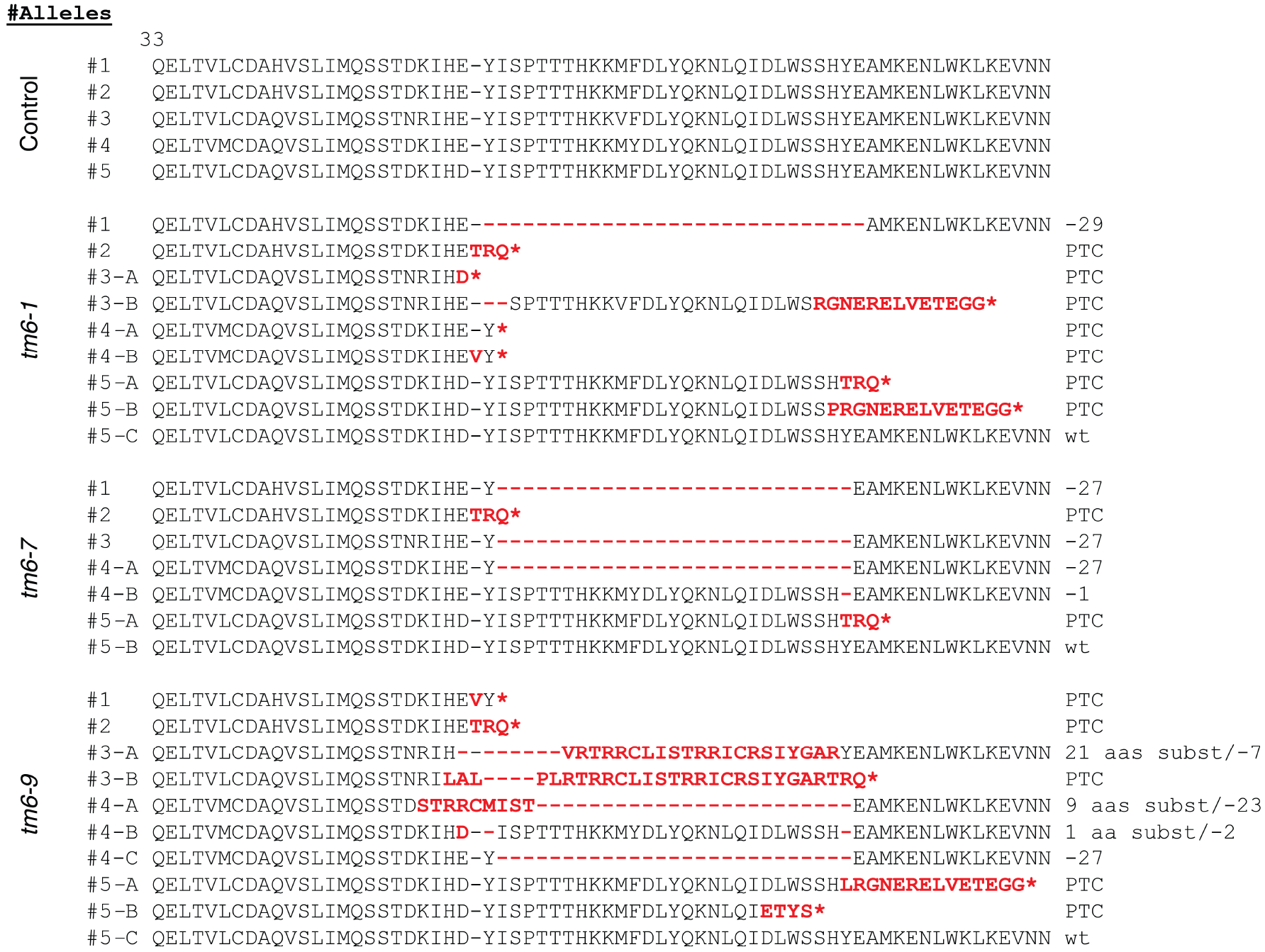
Alignment of TM6 predicted amino acid sequences. TM6 protein sequence from amino acid 33 to 99 in control is aligned with the protein sequences of the *tm6* mutant lines. Red and bold fonts indicate CRISPR/Cas9-induced variants. Red asterisk: premature termination codon (PTC). Information about the amino acid modification is included after the protein sequence.

**Supplemental Figure 6.**
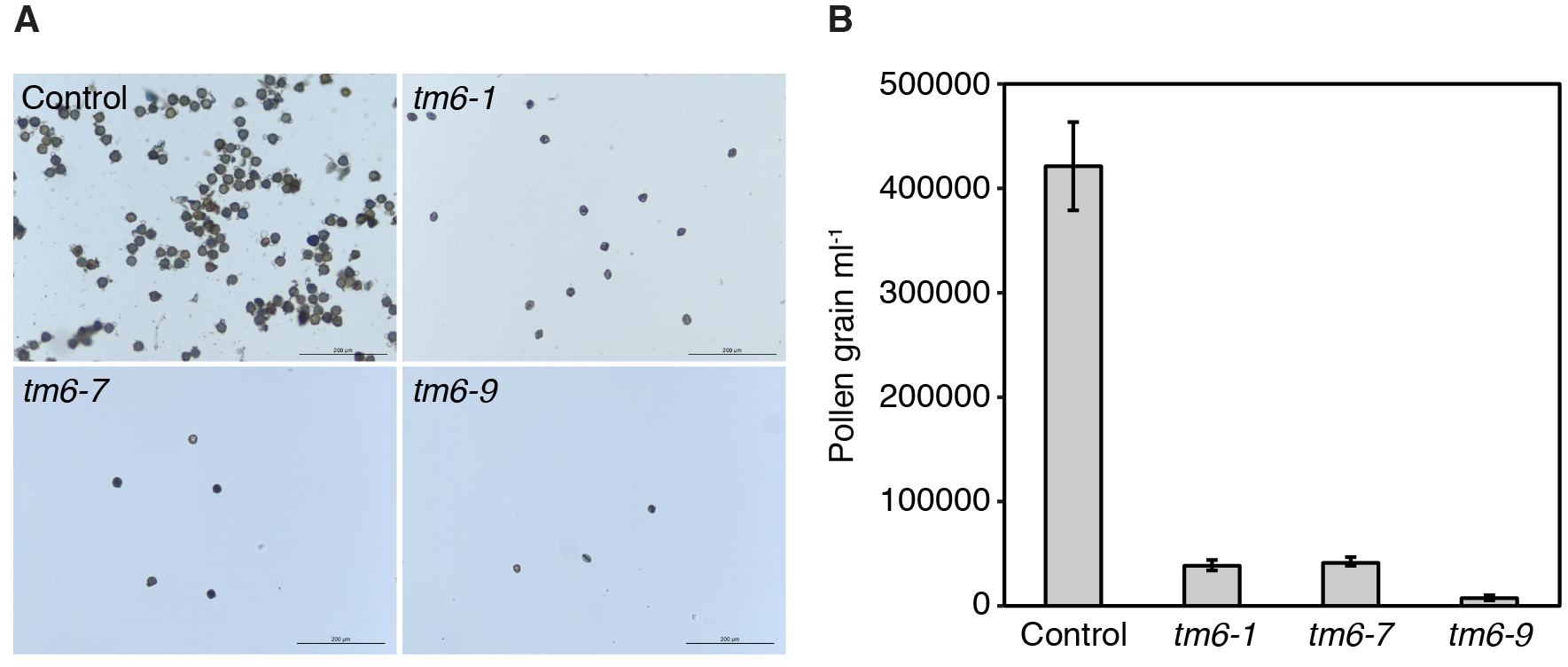
Pollen yield quantification. (**A**) Pictures of pollen grains stained with acetocarmine. (**B**) Quantification of pollen amount using the Neubauer chamber. Error bars denote the standard deviation (s.d.) of three biological replicates.

**Supplemental Figure 7.**
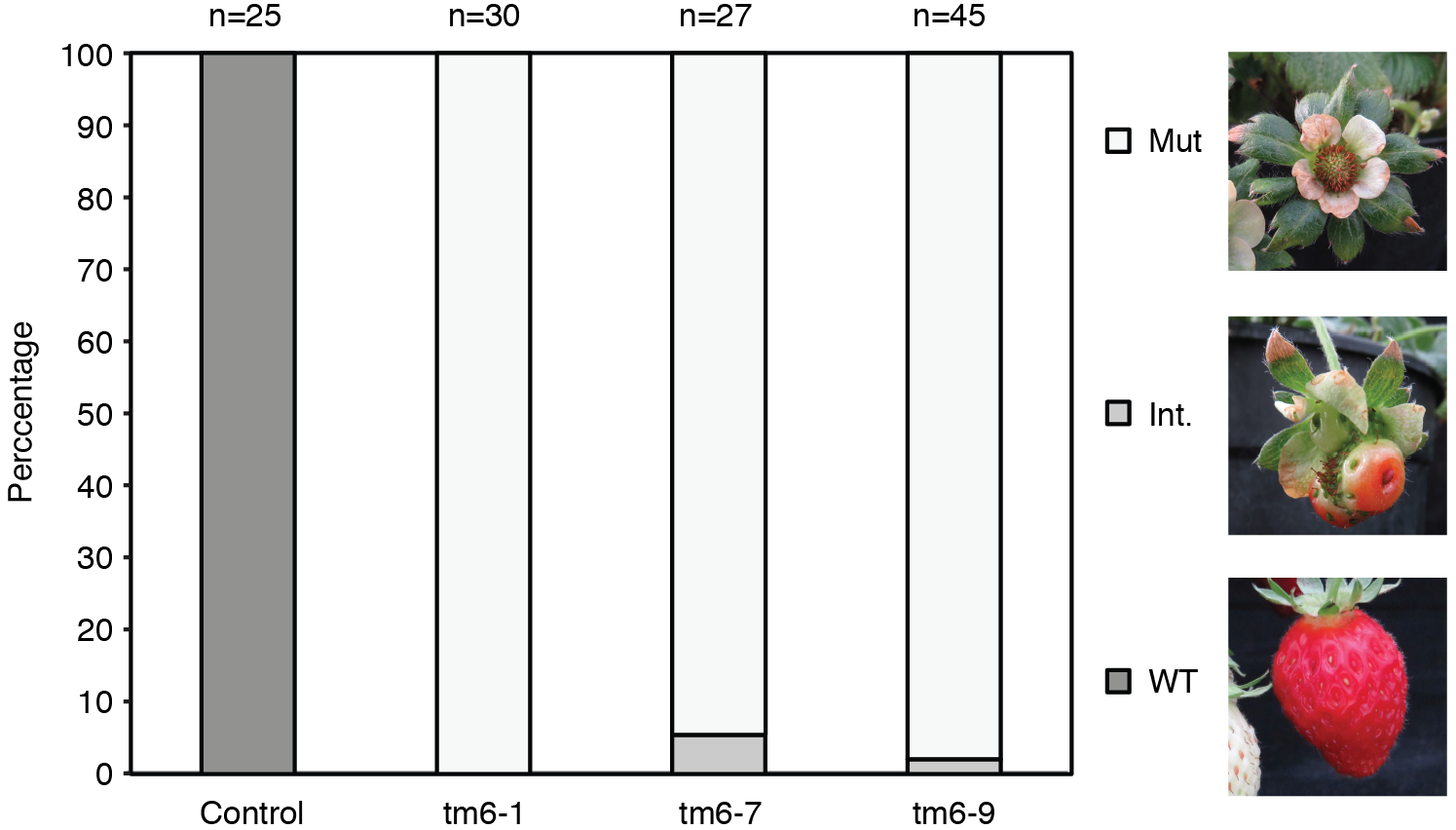
Fruit phenotype quantification. Chart showing the percentage of fruits with mutant, intermediate (Int.) and *wild-type* phenotype in control and *tm6* lines. Fruits with partial receptacle enlargement were considered to have an intermediate phenotype. Numbers of fruits analyzed for each genotype are indicated above the bars.

**Supplemental Table 1.**
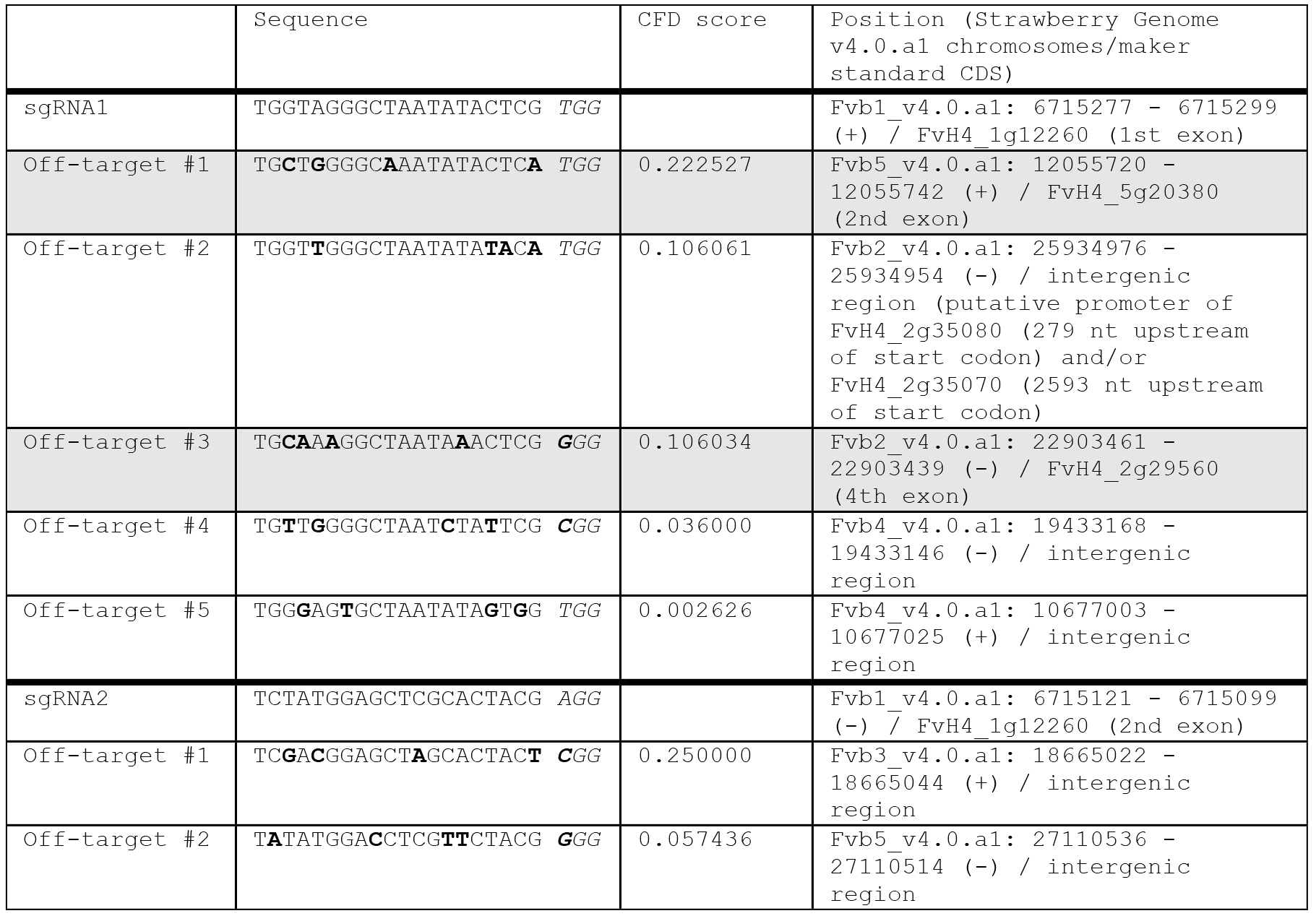
Off-target analysis for sgRNA1 and sgRNA2. Sequences, Cutting Frequency Determination (CFD) score (Doench et al., 2016), and position in the *F. vesca* v4.0.a1 reference genome (Edger et al., 2018) is displayed. CFD score are predictive of off target potential of sgRNA:DNA interactions. Off-targets are ranked by CFD off-target score from most to least likely. Mismatches compared with the sgRNA sequence are shown in bold type. Off-targets located within coding sequences (CDS) are marked in grey.

**Supplemental Table 2.**
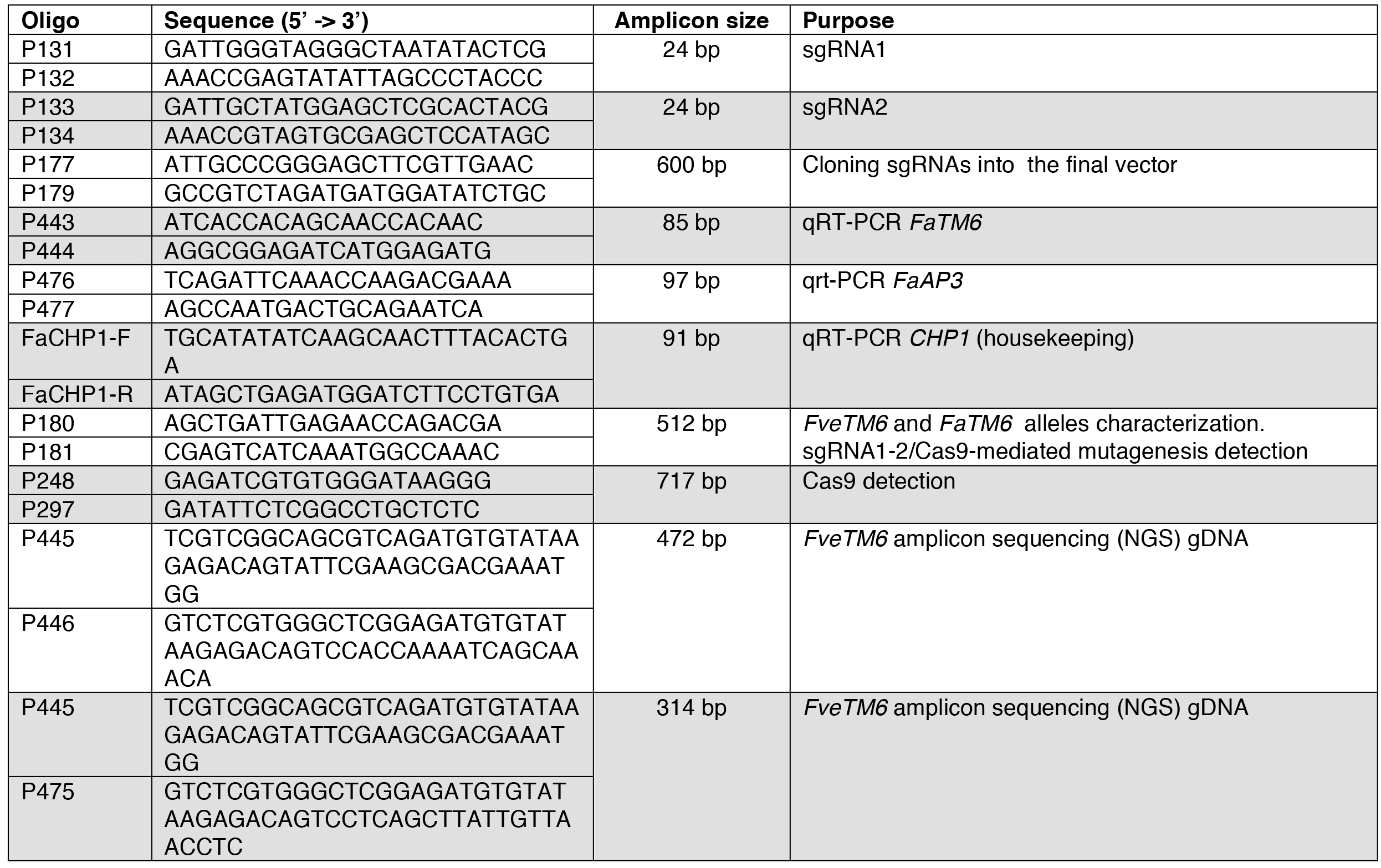
List of oligonucleotides used in this study.

**Supplemental Table 3. Analysis of high-throughput sequencing of amplicons of TM6 cDNA and genomic DNA.***TM6* sequences flanking the two target sites were obtained from cDNA from petals and stamens of *F*. × *ananassa* cv. Camarosa (sheets 1 and 2), and from gDNA from leaves of control and *Fatm6* lines (sheets 3-6). % Prevalence indicates the presence of each cluster obtained by *de-novo* assembly.

